# Comparison of polygenic risk scores for coronary artery disease highlights obstacles to overcome for clinical use

**DOI:** 10.1101/2020.08.09.243287

**Authors:** Holly Trochet, Justin Pelletier, Rafik Tadros, Julie G Hussin

## Abstract

Polygenic risk scores, or PRS, are a tool to estimate individuals’ liabilities to a disease or trait measurement based solely on genetic information. One commonly discussed potential use is in the clinic to identify people who are at greater risk of developing a disease. In this paper, we compare three PRS models that incorporate a large number of genetic markers for coronary artery disease (CAD). In the UK Biobank, the cohort which was used at some point in the creation or validation of each score, we calculated the association between CAD, the scores, and population structure for the white British subset. After adjusting for geographic and socioeconomic factors, CAD was not associated with the first principal components of genetic diversity, which reflect fine-scale population structure. In contrast, all three scores were confounded by these genetic components, highlighting that PRS may be influenced by genetic factors not directly causal for CAD, thereby raising concerns about their biases in clinical application.Furthermore, we investigated the differences in risk stratification using four different UK Biobank assessment centers as separate cohorts, and tested how missing genetic data affected risk stratification through simulation. We show that missing data impact classification for extreme individuals for high- and low-risk, and quantiles of risk are sensitive to individual-level genotype missingness. Distributions of scores varied between assessment centers, revealing that thresholding based on quantiles can be problematic for consistency across centers and populations. Based on these results, we discuss potential avenues of improvements of PRS methodologies for usage in clinical practice.

## Introduction

In the last decades, the need for improvement of risk prediction for human disease has led to the development of many genetic risk scores, which are metrics that use solely genetic data to predict the likelihood of an individual to develop disease. In this highly active area of research, most papers cite the potential clinical value of genetic risk scores in identifying individuals at high risk of developing the disease for early intervention [1]. With the proliferation of direct-to-consumer genotyping panels and as the cost of whole genome sequencing continues to fall, we may not be far from the day where one’s genome sequence becomes a standard part of one’s medical record.

In anticipation of this, a number of polygenic risk scores (PRS), which are the subset of genetic risk scores that explicitly use a large number of genetic markers (thousands or millions) from across the genome, have been published for many diseases, with coronary artery disease (CAD) seeming to be a popular and well-respected choice for potential clinical use [2, 3]. While PRS models can incorporate variants based on various significance thresholds, most scores include variants that do not reach genome-wide significance, and which have not been consistently replicated and functionally validated. Therefore, these may not have well-established associations with CAD. PRS are also used more and more as tools in research studies to help uncover links between traits and mechanisms of disease susceptibility. For instance, they have been used as the genetic instruments in Mendelian randomization studies to establish the causal relationship between an exposure and an outcome [4, 5]. They also have a potential clinical application—namely the stratification of individuals according to their risk of disease as predicted by their genetics, allowing for those at high risk to be monitored more closely or to be given medical interventions before the onset of the disease [6, 1]. Guidelines are now being put forward [7] for appropriate clinical application and reporting of these scores.

Population structure is a concern in medical and statistical genetics, as it may lead to spurious results in association studies, and PRS inherit this problem. It has been previously established that health outcomes in the UK Biobank (UKB) are associated with the population geographic distribution, as are their corresponding PRS [8]. It has also been shown that PRS can be biased by recent demographic history and environmental structure that cannot be corrected for using principal components (PCs) [9]. Additionally, differences in technologies (sequencing, genotyping), quality control (QC) pipelines, imputation pipelines and need for lift over from one genome assembly to another, are all sources of potential missing data in genetic data sets. This may result in the removal of markers that are defined in the score, especially of low-frequency variants [10]. Because these QC steps are performed within a cohort, markers that are kept will vary between cohorts.

PRS are often assessed by testing how well they predict the phenotype (with or without other covariates in the model) or by dividing individuals according to quantiles, with the lowest and highest quantiles being of particular interest. They are also usually normalized within the cohort to assess the mean effect on risk per standard deviation increase in the score. These scores are generally touted for their ability to identify “high-risk” individuals for early intervention, however it is generally hard to define what the appropriate PRS threshold would be in order to determine who these high-risk people are. Therefore, important questions about data completeness, population structure, and appropriate threshold definition arise, highlighting the need for careful examination of what exactly is captured by the proposed scores, before their translation into clinical care can become a reality.

In this study, we examine three recent and well-cited scores for coronary artery disease (CAD) [3, 2, 11]. All three scores tuned parameters in subsets of the UKB, and were then validated on the rest of the cohort. The development approaches varied: the metaGRS is defined on 1,745,179 markers and combined weights from previously published CAD PRSs, the K2018 used the LDpred algorithm [12] on 6,630,150 markers, and the E2020 was constructed on 40,079 markers using the lassosum software [13]. Since all three scores were built using the UKB [14], we revisited these scores in this cohort to investigate three important aspects for which any genetic risk score should demonstrate robustness before it can be successfully implemented for clinical use: population structure, absolute thresholding for high-risk/low-risk individuals, and missing data. We show that all three scores demonstrate limitations on the basis of these criteria and explain why one should be careful in applying PRS in clinical practice to identify future outcomes. If these scores are fundamentally biased by both biological and technical features intrinsic to the genetic data they use, then this demands methods that account for the effects we describe. We also discuss potential avenues of improvements for PRS methodologies before they should be cleared for clinical use.

## Results

### Population Structure

Population structure refers to systematic allele frequency differences within and between populations due to various factors such as mutational events, genetic drift, natural selection, and demographic events [15, 16]. Such variations can lead to misleading results in genetic association studies [17, 18, 19], as genetic markers may appear to be associated with a trait when in fact, they are not biologically related to it. One way to examine population structure is through principal component analysis (PCA) [20, 21], which visualizes the distance and relatedness between populations/individuals. While PRS and PCs are derived from different methodologies with distinct objectives, they both provide weighted summaries of an individual’s genetic markers.

#### Fine-scale Population Structure in the White British

By far the largest group in the UKB are those who self-identified as “British” on the ethnicity question (UKB field 21000) and who clustered together in the PCA provided by the UKB. These individuals, henceforth known as the “white British”, comprise 81.45% of the data set. We performed a principal component analysis specific to this group (Figure 1A and 1B). We see some separation of those born in Wales from those born in Scotland/Ireland. The participants who were born in England are the overwhelming majority of the cohort, and they appear throughout the PCA plots. We also plotted individuals based on the geographic coordinates of their place of birth within the UK (specifically on the island Great Britain) and colored them according to their measurements on the first three PCs (Figure 1C). This showed differentiation between the three nations of Great Britain (England, Wales, and Ireland) and evidence of population clines in England and Scotland.

**Fig. 1.**
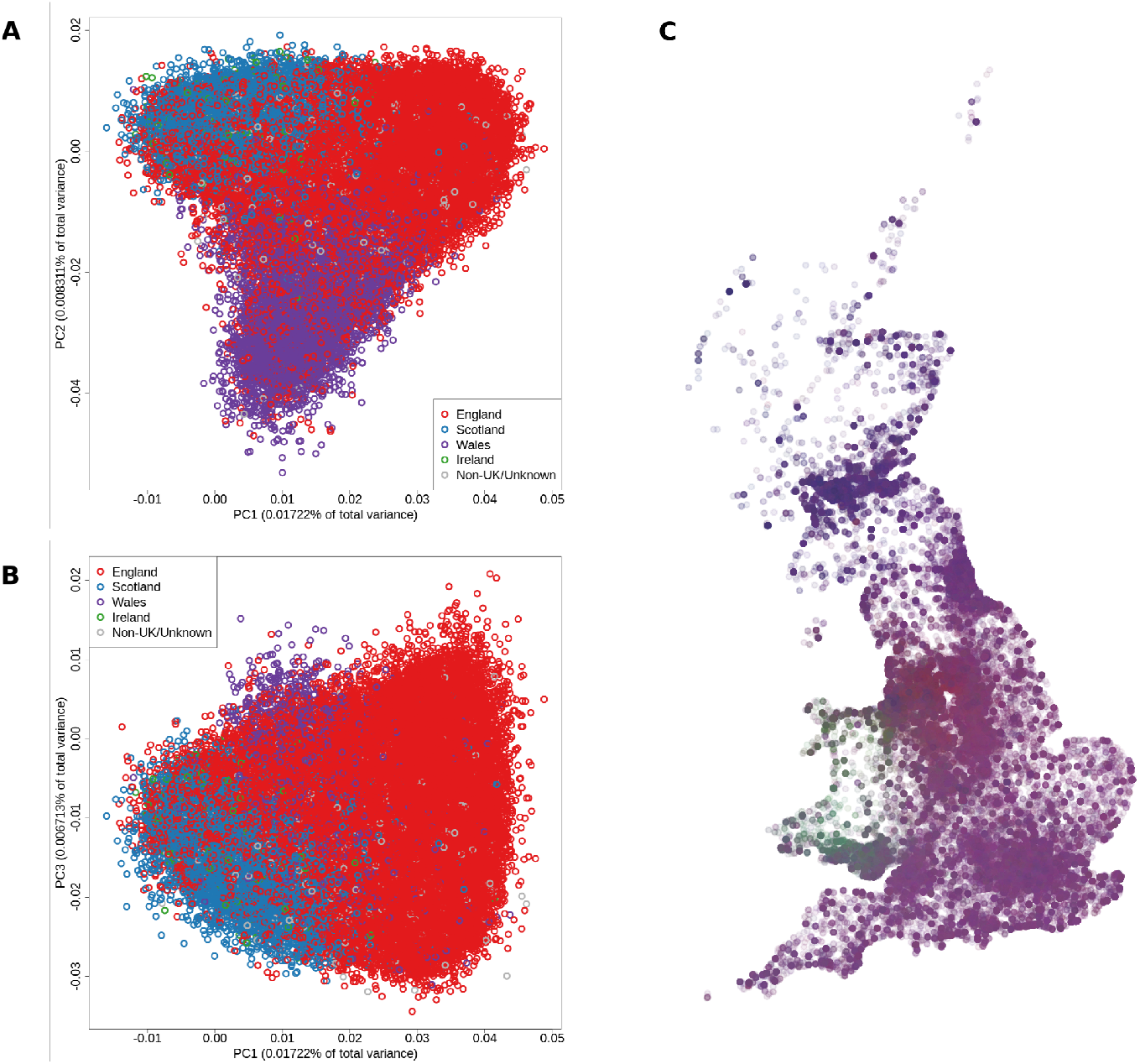
Visualization of principal component analysis (PCA). **A**: First two principal components of the PCA conducted in the white British subset of the UK Biobank, where individuals are colored by their country of birth (England, Scotland, Wales, Ireland, or unknown). **B**: First and third principal components of the PCA conducted in the white British subset of the UK Biobank, where individuals are colored by their country of birth (England, Scotland, Wales, Ireland, or unknown). **C**: White British individuals plotted according to their birthplace in the UK, colored by the first three PC values. Redder values correspond to a higher PC1 value compared to the other two, bluer values correspond to a higher PC2 value compared to the other two, and greener values correspond to a higher PC3 value compared to the other two. Individual dots are slightly transparent to aid in the visualization of the density of people for each place at birth location.

Unsurprisingly, the first three PCs are all strongly associated with the geographic distribution of individuals, both at birth and at the time of assessment (see Methods, Section Covariates used in regression analyses). Both the home location at assessment and place of birth variables were associated with each PC at a level of *p <* 10^*−*660^ for all univariate associations and for multivariate associations fitting all three PCs at once.

We computed three published PRS for CAD: the metaGRS by Inouye *et al*.[3], the K2018 score by Khera *et al*. [2], and the E2020 score by Elliott *et al*.[11] (see Methods for details on building each score). While these methodologies conventionally correct for population structure at the single variant level, these corrections may not fully negate biases when variants are aggregated into a PRS. Indeed, we found that the PCs were associated with each PRS (Table S1). One might interpret a relationship between a PRS and a PC as reflecting true differences in disease risk across the population, but this doesn’t imply a straightforward association between CAD and genetic structure. Several CAD risk factors, such as age, smoking behavior, and indices of deprivation, are also associated with the first three PCs and could be responsible for association with the disease. Additionally, CAD prevalence shows an association with individuals’ places of birth, as well as their home locations, which are also associated with the first three PCs. Therefore, we interrogated whether the genetic differentiation captured by the PCs is a risk factor for CAD independently of these environmental factors, including geographic coordinates.

We adjusted for age, sex, place of birth, home location, Townsend deprivation index, income, age when completed full time education, and smoking status (Table S2). In all cases, none of the PCs is even nominally associated with CAD itself (Table S3), while we see at least one PC associated with each PRS at a significance level of *p ≤* 1.6 *×* 10^*−*7^. In Figure 2, we plot the *p*-value of association between the PCs and the different risk scores (Panels A, B, and C) under different linear regression models. The simplest model (on the far left in each panel) uses just the PCs, and progressively we add age and sex, and various social, environmental, and geographic variables to the model.To provide further clarity on the direction and magnitude of these associations, we have included the regression coefficients for each PC in the supplementary Table S3. In all cases at least one PC remains strongly associated with each PRS, even after these adjustments. In contrast, in Panel D, we perform the same analysis, but as a logistic regression of CAD on these same models, again starting with just PCs and progressively adding more covariates. In this case, once all of these variables are added to the model, none of the PCs is even nominally associated with CAD. These results are also reported in Table S4, along with the Akaike information criterion, which shows that the addition of social and environmental covariates improves the model fit. Therefore, the direct associations we observe between the PRS and the PCs (Table S1) do not reflect a true population cline of CAD risk along the PC axes. These results indicate that the scores capture differences in susceptibility to CAD within the white British population due to the correlation between population structure and environmental risk factors for the disease, rather than differences in the frequencies of alleles that directly affect genetic predisposition.

**Fig. 2.**
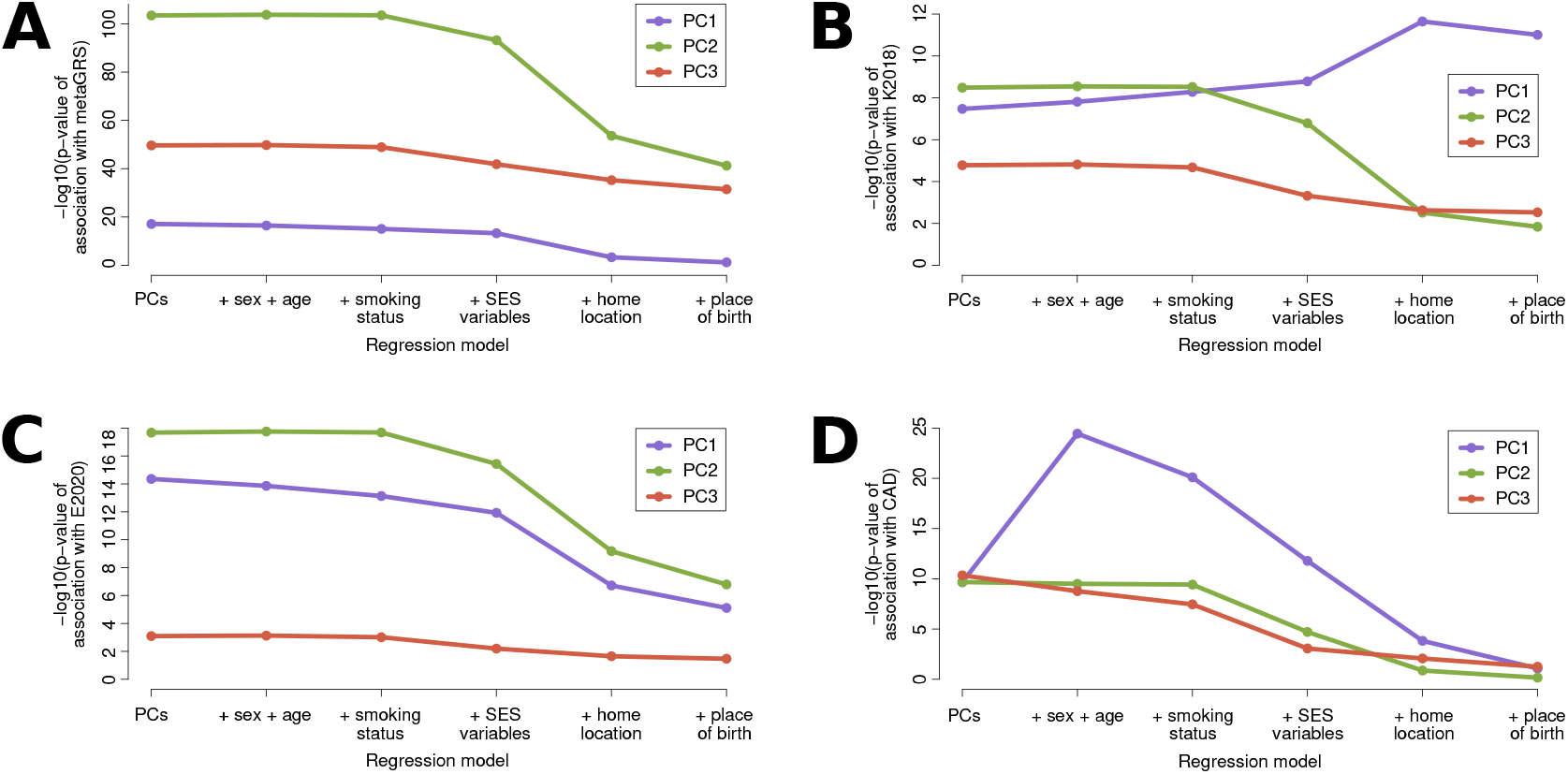
Significance of association between specific PCs (depicted by color) and **A)** the metaGRS, **B)** the K2018 score, **C)** the E2020 score, and **D)** CAD. *−* log_10_ (*P* -values) are shown for different regression models, each successively adding covariates to the one that preceded it. The first model regressed the score or CAD on just the first three PCs; the second included the first three PCs and added sex and age; the third added smoking status (current, previous, or never); the fourth added socioeconomic variables (Townsend deprivation index, income, their interaction effect, and age completed education, adapted to include university); the fifth added home location at assessment (northern and eastern coordinates); and the sixth model added place of birth in the UK (northern and eastern coordinates). All regressions were performed on the same 334,181 white British individuals who had no missing data on any of the potential covariates.

#### Risk Scores in Subpopulations of the UK Biobank

To date, most genetic research has been conducted in individuals of European ancestry, and so most genetic risk scores have been built based on data from European populations. For the three scores under investigation in our study, we observe that non-European subpopulations of the UKB (South Asian; Black British, African, or Caribbean; and Chinese) show different distributions of each CAD risk score compared to the European ones (white British, Irish, and other white) (Figure 3). However, the pattern is not the same for all scores. The raw values for the Black, African, and Caribbean group (here after referred to as the “African” group) and the South Asian group are centered close to zero for the metaGRS, with the distributions of values from the European and Chinese groups shifted to the left and right, respectively. For both the K2018 and the E2020 scores, the distribution of scores from African individuals is the most different compared to the others, with the mean clearly shifted to the left of the other populations. For both of these PRS, the distribution of the scores in people of Chinese descent is similar to that of Europeans, but appears slightly shifted towards the right, further away from the African distribution. South Asians’ distributions, however, overlap well with that of Europeans in both cases, in line with the closer genetic relationship between Europeans and South Asians, which contrast sharply with what is observed with the metaGRS. Thus, depending on the score, the differences in distributions may or may not be aligned with the level of differentiation between these populations. We note that these shifts in distributions do not follow the pattern of prevalence of CAD for each of these groups (Table S5). This raises intriguing questions about the degree to which CAD is driven by the same genetic factors in each of these populations, which merits in-depth investigation in future studies.

**Fig. 3.**
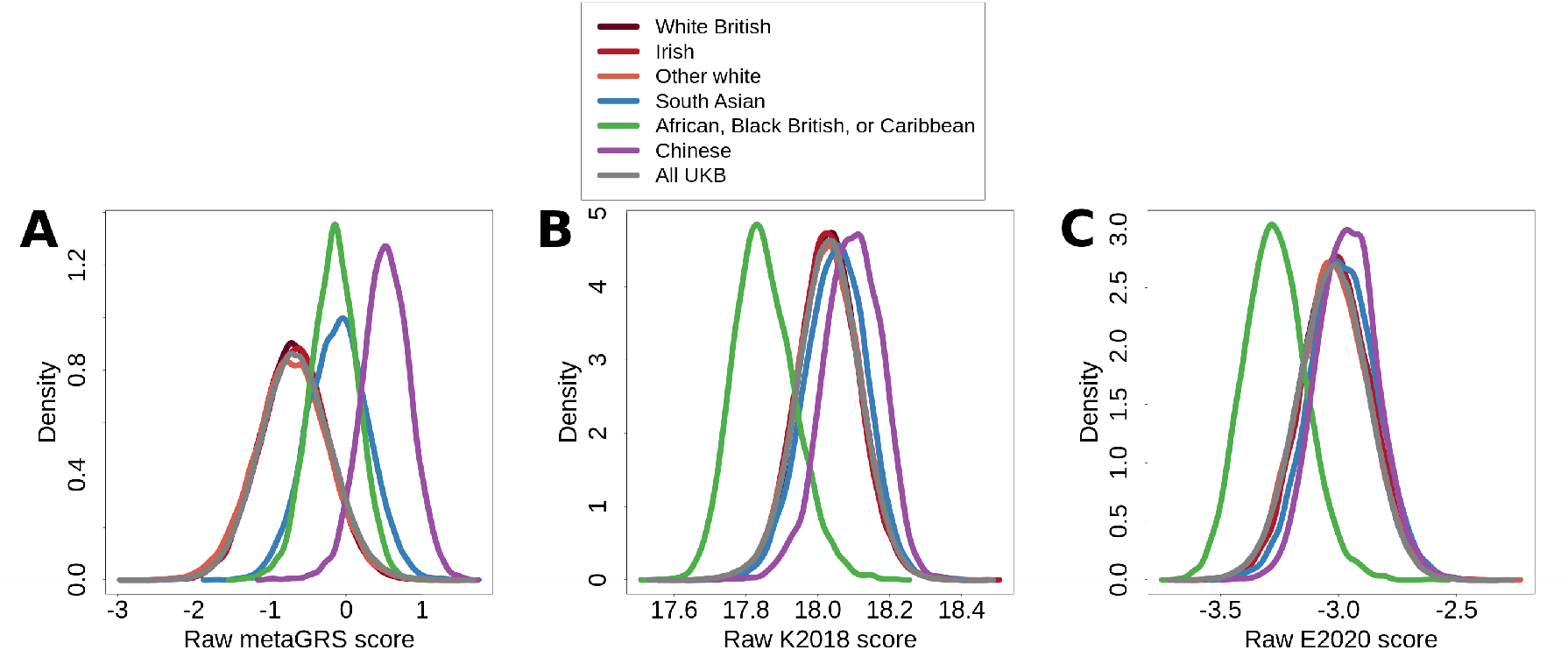
Distributions of raw risk scores of the three risk scores for different sub-populations of the UK Biobank (UKB) : **(A)** metaGRS, **(B)** K2018 score, and **(C)** E2020 score.

Furthermore, it has been demonstrated that PRS have much lower correlations with their target traits in African and East Asian populations than they do in Europeans [22, 23]. We tested if these findings are replicated for the PRS under investigation. While the non-European subpopulations (especially the Chinese) are underpowered compared to the European ones, the point estimates for the effect sizes of each score are consistently lower for the non-Europeans than for the Europeans, except for the E2020 score, where South Asians have a higher point estimate and lower *p*-value than for the “other white” group (Table S5). Additionally, the estimated effect sizes on the scores are considerably lower in the African group than in all the others.

### Thresholds for Risk

PRS are usually evaluated within a cohort, by dividing individuals according to quantiles of their PRS, with the lowest and highest quantiles being of particular interest. The ability to identify “high-risk” individuals for early intervention is often used as a selling point of these scores, although there is often no indication given on what an appropriate threshold would be for the PRS in order to determine who these high-risk people are. Furthermore, if scores vary across the population in relation to population structure and geography, each institution will have a unique mean and standard error. To test this, we calculated the means and standard deviations for each score in white British individuals who attended four different UKB assessment centers, and for the white British cohort as a whole (Table 1). Additionally, we calculated the empirical thresholds for the top 10 and 5% of risk scores at each center. Each center has a subtly different distribution of each of the scores from the others, whose statistical significance we tested using Kolmogorov-Smirnov tests between each pair of centers for all three scores. We indicate the ones that were significantly different (*p <* 2.78 *×* 10^*−*3^, to account for multiple testing) in Table 1 and report the *p*-values of these tests in Tables S6, S7, and S8.

**Table 1.** Sample size, mean, standard deviation (SD), and thresholds for the top 10% and top 5% of risk scores among white British participants from four different UK Biobank assessment centers, as well as the whole subset (“All”). For each center and each score, we calculated the mean and SD for the subset of white British participants who attended a given assessment center. We also calculated these values for the full cohort. We report significant difference of distributions (*P*-values of Kolmogorov-Smirnov tests *<* 0.00287, which is the Bonferroni-corrected 0.05 threshold) compared to Reading (^*r*^), Cardiff (^*c*^), Newcastle (^*n*^) and Glasgow (^*g*^) assessment centers. All *P*-values of Kolmogorov-Smirnov tests of the pairwise equivalence of the distributions between assessment centers are reported in Tables S6, S7 and S8.

| Score |  | Reading | Cardiff | Newcastle | Glasgow | All |
| --- | --- | --- | --- | --- | --- | --- |
|  | Size | 22,635 | 15,462 | 28,053 | 14,205 | 408,567 |
| metaGRS | Mean | $^{n,g}$-0.7127 | $^{n,g}$-0.7115 | $^{r,c}$-0.6675 | $^{r,c}$-0.6617 | -0.6907 |
|  | SD | 0.4478 | 0.4501 | 0.4474 | 0.4479 | 0.4483 |
|  | 10% | -0.1380 | -0.1329 | -0.0893 | -0.0871 | -0.1153 |
|  | 5% | 0.0257 | 0.0453 | 0.0724 | 0.0753 | 0.0500 |
| K2018 | Mean | $^n$18.0323 | 18.0332 | $^r$18.0363 | 18.0330 | 18.0340 |
|  | SD | 0.0850 | 0.0856 | 0.0844 | 0.0848 | 0.0849 |
|  | 10% | 18.1406 | 18.1432 | 18.1456 | 18.1407 | 18.1427 |
|  | 5% | 18.1719 | 18.1747 | 18.1772 | 18.1736 | 18.1746 |
| E2020 | Mean | $^{n,g}$-3.0219 | $^g$-3.0186 | $^n$-3.0152 | $^{r,c}$-3.0138 | -3.0193 |
|  | SD | 0.1453 | 0.1446 | 0.1450 | 0.1456 | 0.1451 |
|  | 10% | -2.8346 | -2.8332 | -2.8300 | -2.8264 | -2.8330 |
|  | 5% | -2.7843 | -2.7763 | -2.7773 | -2.7739 | -2.7806 |

Where differences in the distributions of the scores exist, they appear between the two northern cities (Glasgow and Newcastle) and the two southern ones (Cardiff and Reading). The metaGRS shows the strongest and most consistent differences between these groups. In contrast, the K2018 score has the most consistent distributions across the four centers, with a statistically significant difference between Reading and Newcastle only, which may be detectable due to the relatively larger sample sizes in these two centers. Finally, the E2020 score shows the strongest differences between Reading and the two northern cities, though there is also a statistically significant difference between Glasgow and Cardiff.

#### Concordance of the Scores

Because PRS tend to be defined in relative terms, one natural question when looking across multiple ones is how consistent the results are—that is, given a cohort, do different scores identify the same individuals as high-risk? To answer this question, we calculated the Spearman rank correlation between each of the three pairs of CAD PRS investigated here in the white British subset of the UKB. The highest correlation is between the metaGRS and the K2018 scores, which is 0.7676. The next highest was between the K2018 and the E2020 scores, at 0.6713. Finally, the correlation between the metaGRS and the E2020 scores was 0.5797. These correlations are not as high as one might expect, suggesting that there is a good deal of variation among the scores in who will be identified as high-or low-risk.

We next looked at the high- and low-risk individuals for each score to evaluate their overlap. We report the proportion of overlap of white British UKB participants who scored in the highest and lowest 10% of each score (Table S9). In the best case scenario, when comparing the metaGRS to the K2018 score, a little over half (55.30%) of individuals are identified as being in the top 10% of risk by both scores. The proportion of overlap among all three scores was 0.2917 for the low-risk group and 0.2912 for the high-risk one.

While there is some consistency across the scores in who is identified as high-or low-risk, the agreement among the scores is not as strong as we might expect or want for individuals in the extremes of the distribution. Additionally, there are a small number individuals who are defined by one score as being high-risk and by another as low-risk (highest or lowest decile of risk score), reported in Table S10. Here again, the largest discrepancies were observed between the metaGRS and the E2020 scores, where a total of 195 individuals were placed in the opposite extreme risk category by the other two scores. These analyses raise questions about which score is the most accurate or appropriate in a clinical setting; however assessing this is beyond the scope of this paper.

### Impact of Missing Data

Missingness in genotypic data poses a significant challenge in genomic studies. Most genetic analyses typically tolerate a missingness threshold of around 5%, with some researchers adopting a more stringent criterion of 1%. However, as PRS become increasingly discussed in clinical and research settings, the implications of missingness on these scores remain under-investigated. Genotyping chips, imputation pipelines, and quality control filters differ among cohorts, which may cause difficulties in calculating PRS due to the removal of markers that are defined in the score, thereby creating inconsistencies in genotyping data across cohorts, even when they use the same technology. In addition to cohort-wide missing genotype data, most QC pipelines tolerate some amount of missing genotype data for each individual. This means that most, if not all, participants are missing genotypes at a small set of random markers, which changes from person to person. To handle this type of missingness when calculating the risk scores in each cohort, the mean effect allele dosage calculated in the rest of the cohort is often used. This discounts the effect of the marker on the score of that individual, but has downstream consequences on the interpretation of individuals’ levels of risk. Presently, the field lacks standardized benchmarks for acceptable missingness rates in PRS studies, creating the risk of heterogeneous practices and potential inconsistencies across different clinical settings.

To investigate the effect of individual missingness, for each PRS, we identified the 200 people with the most extreme scores (the highest 100 and the lowest 100). We then randomly removed their genotypes at markers used in the scores. We repeated this process 10 times and took the mean of these new scores over all replicates for these individuals (see Methods Section Simulating missing genotype data in the UK Biobank) and report how many individuals still had the 100 highest or 100 lowest scores for each level of missingness (Table 2)

**Table 2.** Number of people who originally had the highest and lowest 100 scores for each risk score who, after setting a random group of genotypes missing, still have the most extreme risk scores.

| Missingness | Extreme | metaGRS | K2018 | E2020 |
| --- | --- | --- | --- | --- |
| 0.01 | Highest | 89 | 92 | 91 |
|  | Lowest | 96 | 94 | 91 |
| 0.02 | Highest | 82 | 87 | 85 |
|  | Lowest | 90 | 85 | 86 |
| 0.05 | Highest | 60 | 69 | 64 |
|  | Lowest | 71 | 70 | 66 |
| 0.10 | Highest | 35 | 49 | 39 |
|  | Lowest | 45 | 45 | 35 |

Predictably, as individual missingness increases, the number of people whose scores remain in the extremes drops. In practice, it is unlikely that anyone with a genotype missing rate as high as 10% would be retained in analysis, but it is useful to see the trajectory. The reason for this phenomenon is illustrated in Figure S1, which shows that for each score, as missingness increases, the scores tend towards the mean.

When allele frequency data from a wider cohort is not available, as it would be the case when computing a PRS for a specific patient in the clinic, there is no way to calculate a mean effect allele dosage. In this case, missing markers are effectively removed from the score, which is the equivalent of treating individuals as though they are homozygous for the non-effect allele at the missing marker. For scores that are defined solely in terms of risk increasing alleles (as is the case with the K2018 score) this will result in individuals with high missing genotype rates having lower risk estimates than those whose missing genotype rates are lower. When the score is defined as a mix of risk increasing and decreasing alleles (as the metaGRS and E2020 scores are) then individuals with higher missingness rates will tend to have less extreme scores. These results demonstrate that individual variation in missingness rates will impact the stratification of disease risk in the cohort.

## Discussion

In this study, our first aim was to elucidate the complex interplay between genetic factors, population structure, and disease risk prediction in the case of CAD, for which multiple PRS have been developed and shown to capture genetic risk. We highlighted that all three PRS under investigation predict statistically significant differences across the white British population on the basis of population structure captured by PCs, even after adjustment for environmental variables. However, these PCs do not appear to be associated with the target trait, CAD, once these same environmental variables are accounted for. It also appears that the models that account for environmental effects fit the data better, according the AIC (Table S4). This suggests that the differences in genetic risk across the population, predicted by the scores, are spurious. This association between the scores and genetic PCs will result in the misclassification of individuals’ genetic risk, as those whose genetic PCs are in the extremes of the distributions are more likely to have scores that put them in the high-or low-risk category according to the PRS. In a clinical setting, this could lead to unintentional and erroneous pathologizing of genetic ancestry. Our primary concern, particularly in the context of PRS utility in clinical settings, is the existence of such an effect in the first place, rather than its magnitude. From a clinical perspective, even a minimal effect that persists after rigorous statistical control can have implications for the robustness and reliability of PRS scores, especially when population structure comes into play. Therefore, our emphasis was on the existence of these associations and their potential implications for PRS utility.

Furthermore, we detected statistically significant differences in score distributions for individuals from different UKB assessment centers. These differences are problematic because a person’s genetic make-up is constant and so their PRS and the subsequent characterization of their genetic risk ought to be as well: a person’s genetic predisposition does not change when they move to a new city. Additionally, normalizing within the available data means that individuals who are assessed at institutions where people are disproportionately high-risk may miss out on interventions that could be helpful to them. Conversely, people who are assessed at institutions where people are disproportionately low-risk may be given medical interventions inappropriately, exposing themselves to side-effects for minimal potential benefit. We also reported a lower-than-expected concordance between the three PRS studied, and we identified patients that are put in the top 10% of risk by some PRS and the bottom 10% of risk by other PRS. It would be valuable to see if such discrepancies remain when other clinical variables are considered.

As with most other studies of PRS, we have the limitation of neglecting gene *×* gene and gene *×* environment interactions. While one’s genetic make-up does not change when one moves from one city to another, the interaction between one’s environment and one’s genetics might. Mostafavi *et al*. have previously demonstrated that PRS have varying predictive values across various social and environmental strata [24], which suggests that there may indeed be gene *×* environment effects that alter the interpretation of scores across different environments, but we have not investigated this in this work. Furthermore, in order to retain as many samples as possible, we were constrained in our choice of the environmental covariates to use in our models and were not able to choose the ones that were best for capturing these effects (see Methods, section *Covariates used in regression analyses*, for further discussion).

To move the field forward, it is vital that PRS are investigated to assess to what degree they are associated with fine-scale genetic population structure in their target populations. Ideally, they would demonstrate that any association between population structure and the PRS is in proportion to the association between population structure and the trait itself. However, in practice it may not be possible to completely separate a markers’ effect on the trait from an effect it has that is mediated through social or environmental factors.

It is also important to consider the option of absolute thresholds for risk, which would bring these scores in line with traditional CAD risk factors, such as measures of obesity (waist-hip ratio and body mass index), blood pressure, and cholesterol levels. All of these have absolute thresholds that differentiate between low, normal and increased risk, even if some of those thresholds are not constant across ethnic groups or the sexes. Furthermore, the current practice of normalizing within a cohort means that the score cannot be compared across cohorts or populations. This introduces a problem raised by Martin *et al*. [23], where PRS can be used only in the majority ethnic group served by the institution, such that members of minority ethnic groups will miss out on the potential benefits, which will create or exacerbate existing healthcare disparities between these groups. Finally, absolute risk thresholds would make PRS more accessible to patients, since it means they could know their raw score and its interpretation the same way they do for measures like LDL cholesterol or BMI. With advances integrating PRS for CAD into the pooled cohort equation (PCE) [25], this strategy will face the same issues of population structure and data processing as with standalone PRS. It is important to ensure that novel tools aimed at improving risk prediction, once implemented in real-world settings, don’t inadvertently lead to biases or perpetuate health disparities. The potential implementation of CAD-PRS within PCE must be approached with caution to guarantee equitable and accurate risk assessments for diverse populations.

Missing data are inevitable, and with growing interest in PRS, particularly in clinical contexts, addressing this issue becomes critical. Even if whole genome sequencing becomes standard, sequencing data will still need to pass through quality control pipelines which will inevitably lead to a loss of data at the cohort level, as well as for each individual. While a minor level of missing data might be acceptable in broader genomic analyses, even low missingness can skew PRS, potentially leading to inaccurate risk predictions. Given the established thresholds of missingness (e.g., 5%), our results offer insights into the implications of these standards on PRS calculations. There is a need to provide guidance on how to deal with missing genetic data when a PRS is generated. For small genetic risk scores, it may be possible to use proxy markers in LD with the missing one, which are highly correlated with those in the risk score. This becomes more challenging for PRS models that incorporate large numbers of markers, where the best available proxies may also be included in the score. One possible area of future work could be in the development of algorithms that can readjust a score’s weights based on which markers are available. We believe that improvement on current scores is possible. Part of the reason why the problems we have outlined arise is because of the way these scores are constructed and assessed. There is an implicit assumption that people who show a phenotype must have an increased genetic predisposition for it. While this is true for some traits (for example, eye color [26]) there is a large environmental component to CAD which could potentially overwhelm the underlying genetic predisposition. These environmental factors can create cases out of people at low genetic risk for the disease and prevent those at elevated genetic risk from developing it. The process of creating and validating these scores focuses on their ability to predict phenotypes from genotypes. For CAD, this might be a perverse incentive, since it rewards the PRS for including loci whose associations with the trait are mediated through social or environmental covariates—that is, genetic artifacts of social and environmental risk. This means that if two individuals have the same true underlying genetic risk, but one of them develops CAD and the other does not, a potential score that assigns a higher value to the individual who develops CAD will be favored over one that assigns them the same value. One way of avoiding this problem may be to restrict the genetic risk scores only to loci with well-validated associations with CAD. These scores would use a much smaller number of markers, allowing for proxy SNPs to be used when genotype data is missing, and may show less spurious association with population structure.

Finally, it is worth questioning whether building a single PRS for CAD itself is the best way of capturing the genetic liability for this disease, and whether it makes sense to have a single score for both men and women. The interplay between nature and nurture in CAD is complex and multifactorial [27], with both factors influencing disease risk in combination, which is reflected in our analyses. Traditional risk factors were highly associated (*p <* 2 *×* 10^*−*16^ with all three of our scores and a high PRS for any of the scores predicted hypertension, high cholesterol, diabetes, and obesity (measured by BMI and waist-hip ratio), whereas the metaGRS predictive ability was claimed to be mostly independent of established risk factors [3]. Clearly the scores use markers associated with these risk factors, and it may make sense to build separate PRS for each of them, and then combine them into a risk model, as discussed previously [28]. One advantage of these approaches would be that pleiotropic effects could be accounted for. For instance, if a variant increases adiposity, but decreases LDL cholesterol, then these effects can both contribute to the final risk estimation of CAD via the separate scores, whereas in a single score for CAD, one of these effects may mask the other, or they might cancel each other out. Additionally, there might be more scope in using multiple scores to incorporate environmental and non-genetic biological effects, and specifically the effects of age and sex.

In conclusion, as a result of our findings, we propose that PRS should fulfill the following criteria: demonstrate robustness to population structure, provide absolute thresholds for high-risk versus normal or low-risk, and have a way of compensating for missing data. If these criteria are not considered appropriately before these scores are put into clinical use, there is a risk that social problems arising in specific geographic areas, such as poverty and unequal access to quality education, food, and medical care, remain unresolved due to the perception that the groups who suffer disproportionately from these problems are simply genetically more prone to disease.

## Methods

Except where otherwise noted, all analyses were performed in R version 4.0.2. [29]. The code used for the analysis and to produce the figures is available here: https://github.com/HussinLab/trochet_2023.

### Study populations

The UK Biobank (UKB) is a prospective cohort of about half a million individuals from the United Kingdom, recruited between the ages of 40 and 69 [14]. The full dataset is multiethnic, but our analyses were concentrated on the subset of “white British” individuals, defined as those who identified as “British” on the ethnicity question (field 21000) and who clustered together in the UKB principal component analysis (PCA) on PCs 1 and 2, for a total of 408,550 individuals. These people were also identified as “Caucasian” in field 22006 (genetic ethnic grouping). We selected this subset as we wished to avoid confounding due to systemic biases affecting access to and quality of healthcare in the UK [30]. Given that it represents 81.45% of the whole of the UKB, the genetic architecture of a given trait in this population will have a heavy influence on the results of genetic analyses that use the full UKB cohort. We want to emphasize that the exclusion of non-white UKB participants in some analyses presented here does not reflect a lack of interest of studying ethnically diverse human populations. The analyses shown here were conducted under UK Biobank project number 49731, and the project has ethical approval from the Ethics Board of the Montreal Heart Institute, Project 2019-2418.

### Overview of PRS computation

We used three published PRS for CAD: the metaGRS by Inouye *et al*.[3], the K2018 score by Khera *et al*. [2], and the E2020 score by Elliott *et al*.[11], which we computed on all UKB participants, regardless of their genetic ethnic grouping. The subset and the validation set of UKB used to develop the metaGRS and the E2020 scores included all UKB participants, while the K2018 score was restricted to the white British subset (81.45% of the cohort). For the metaGRS, random linkage disequilibrium (LD) pruning was used to generate candidate sets of markers in training data from the UKB (1000 randomly selected prevalent CAD cases and 2000 controls) and the set that had the highest hazard ratio in the training set was selected. Weights were chosen by combining the weights from three previously published CAD PRS plus the effect size estimate of the marker on CAD in the UKB. The K2018 score was constructed using the algorithm LDpred [12], while the E2020 score used the software lassosum [13]. Both pieces of software were used to generate candidate scores, and in both cases, the score with the highest predictive value was selected, measured by area under the curve (AUC) of the receiver operating characteristic (ROC) curve in a logistic regression model with CAD as the outcome.

### Principal component analysis of the white British subset of the UK Biobank

We used flashPCA [31] to calculate the top 50 PCs on the unrelated white British UKB participants, using the imputed genotype data, QCed so that all SNPs had a minor allele frequence (MAF) *≥* 0.01, have genotypes available for at least 99% of samples, a posterior probability of at least 0.9 on the imputed genotype, and whose *p*-values for being out of Hardy-Weinberg equilibrium were *≥* 10^*−*6^. We removed the four regions of high LD/known inversions suggested by the authors of flashPCA and used the `--indep-pairwise` function in Plink v1.9b 5.2 [32, 33] to prune the SNPs using the suggested parameters of a 1000 kilobase window, a step size of 50 variants, and an *r*^2^ of 0.05.

In order to create the subset of unrelated people for the PCA, we removed one individual from each pair of related individuals identified in a file provided by the UKB, yielding 335,088 unrelated participants. We then used the loadings to project all 408,550 white British onto these 50 PCs. We computed the Pearson correlation coefficient between the top 40 principal components provided by the UKB over the whole dataset and our PCs computed on the white British, and found strong correlation between our PC 1 and the UKB’s PC 5 (correlation coefficient -0.961) and between our PC2 and the UKB’s PC 9 (correlation coefficient of 0.917).

### Calculating genetic risk scores

We selected three polygenic risk scores (PRS) from the literature, each predicting the risk of coronary artery disease (CAD) [3, 34, 11]. All three PRS are available at The Polygenic Score (PGS) Catalog [35], where we accessed the necessary information on the SNPs used in the scores, including their respective effect alleles and weights. We downloaded the data contained in this repository and calculated all three scores in Plink v1.9b 5.2 [32, 33] with the `--score` function using the imputed UKB genetic data for each individual from the white British subset.

Unlike the other two scores, a high Elliott score is associated with a decreased risk of CAD. To aid in the comparison across scores, we used the negative of the Elliott score in all our analyses.

### Trait definitions

In the UK Biobank, Coronary artery disease was defined in the same way as it was in Inouye *et al*.’s paper [3], using UKB fields 6150, 20002, and 20004. In the linked medical and death records, we looked for ICD9 codes 410-412, ICD10 codes I21-I24 and I25.2. Among the surgical procedure data, we looked for OPCS-4 codes K40-K46, K49, K50.1, and K75. In the self-reported data, the relevant surgical procedures were recorded as 1087, 1095, and 1581. Unlike the study’s authors, we did not differentiate between incident and prevalent cases. Of the 408,698 white British individuals for whom these data were available, 23,374 (5.72%) met the above criteria for CAD.

### Simulating missing genotype data in the UK Biobank

For each score, we identified the people with the highest 100 and the lowest 100 scores. For each of these individuals, a random selection of genotypes at markers used by the score was set to missing. The score was then recalculated. This process was performed 10 times each for different rates of genotype missingness: 1%, 2%, 5%, and 10%. For each rate, we took the mean score for each individual across all 10 runs as their new score. These new scores replaced the individuals’ original scores, and were renormalized with the rest of the cohort’s original scores.

### Covariates used in regression analyses

We saw in our regression analyses that the *p*-values of association between the PCs and CAD increase further with the addition of variables such as pack years of smoking (field 20161), measures of alcohol consumption, and exercise. However, the inclusion of these variables means the exclusion of increasing numbers of individuals, who are not evenly distributed throughout the dataset with respect to all the relevant variables. For instance, the individuals for whom there is no data on pack years of smoking are disproportionately from the “previous smoker” category. They are also older on average by almost a full year (0.9740) than the group for which these data are available. The variables included in the models in Table S4 were chosen specifically to retain as much of the data as possible and checked to ensure that biases (especially in the distribution of the PCs) were not introduced due to missing data.

It is for this reason that we approximate socioeconomic status using Townsend deprivation index, income, and age when completed full time education. The UK Biobank contains a number of potential measures, including indices of multiple deprivation that were calculated within each of England, Scotland, and Wales (fields 26410, 26427 and 26426, respectively), which were tempting to use, but whose inclusion altered the PC distributions of the remaining sample relative to the original. We report the full list of covariates in Table S2.

## Author contributions statement

HT and JH designed this study. HT performed all analyses with help of JP, who contributed background research on the PRS used. RT provided a clinical perspective for the improvement of these scores. HT and JH wrote the paper together and all authors approved its final version.

## Acknowledgments

We would like to thank Dr Na Cai, Dr Marie-Julie Favé, Dr Ryan Christ, Dr Claude Bhérer, Jean-Christophe Grenier and Pamela Mehanna for their comments on our work. RT and JH are Fonds de la Recherche du Québec en Santé (FRQS) Junior 2 Scholars.

## Supplemental Tables and Figure

**Table S1.**
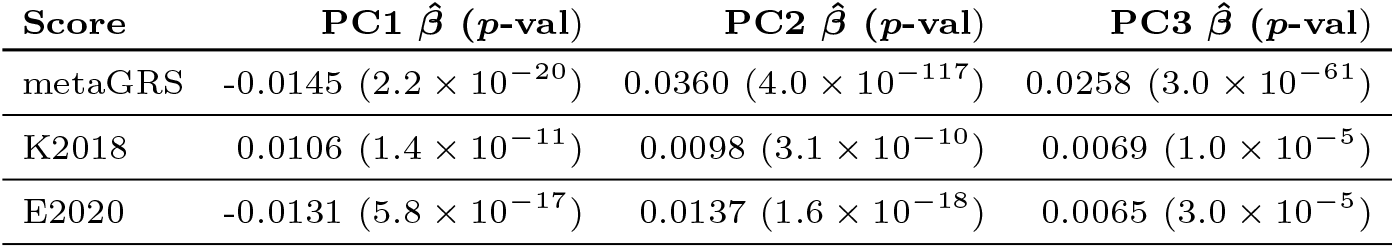
Effect size estimates 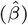 and strength of association (*p*-values) between PCs and scores in the white British subset of the UK Biobank. We provide here the *p*-values of each PC in a linear regression of the score (normalized within the white British subset) on the first three PCs calculated within the white British subset. The PCs were scaled for this comparison.

**Table S2.**
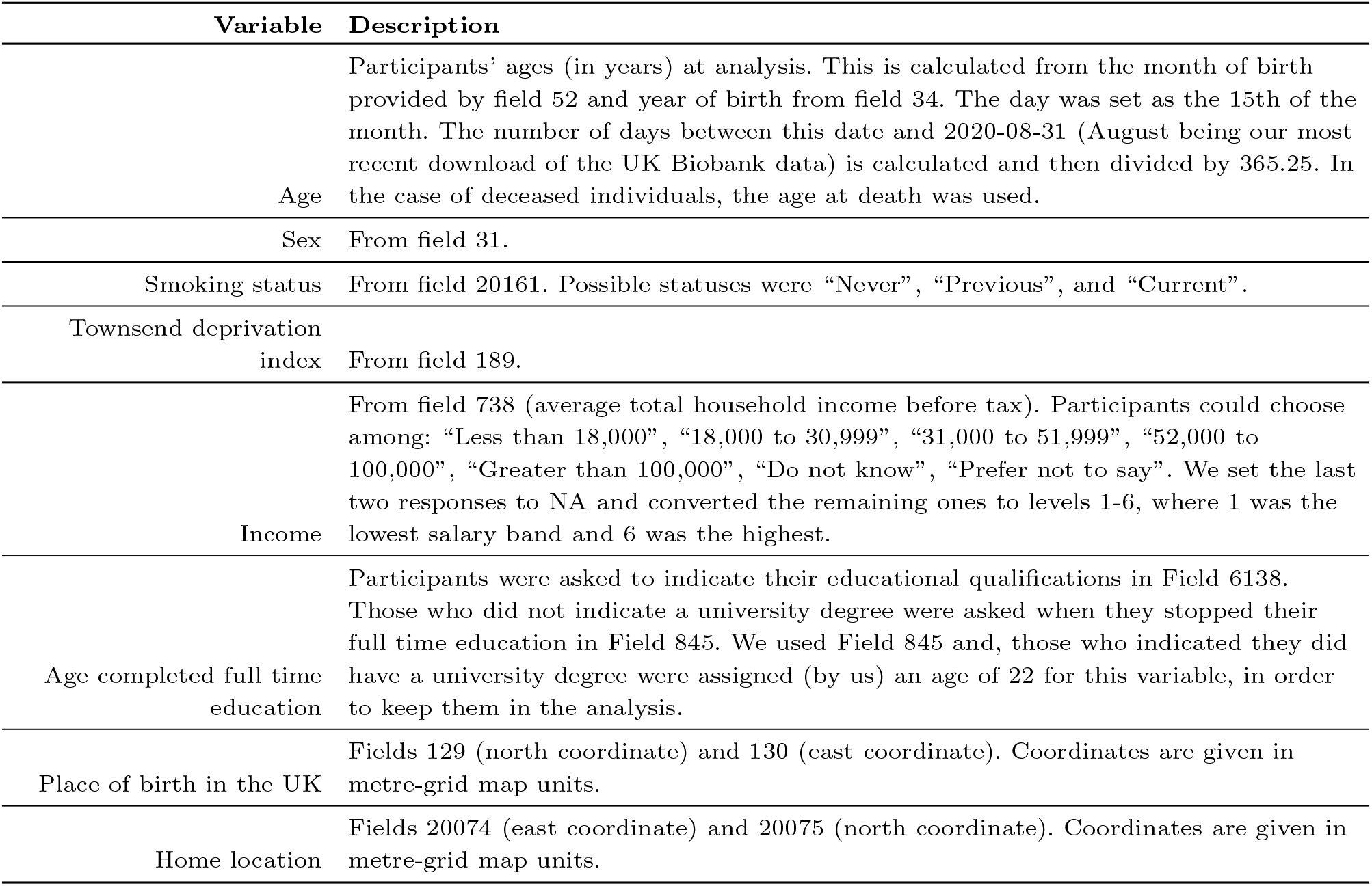
Description of variables used in our regression analyses.

| Variable | Description |
| --- | --- |
| Age | Participants' ages (in years) at analysis. This is calculated from the month of birth provided by field 52 and year of birth from field 34. The day was set as the 15th of the month. The number of days between this date and 2020-08-31 (August being our most recent download of the UK Biobank data) is calculated and then divided by 365.25. In the case of deceased individuals, the age at death was used. |
| Sex | From field 31. |
| Smoking status | From field 20161. Possible statuses were "Never", "Previous", and "Current". |
| Townsend deprivation index | From field 189. |
| Income | From field 738 (average total household income before tax). Participants could choose among: "Less than 18,000", "18,000 to 30,999", "31,000 to 51,999", "52,000 to 100,000", "Greater than 100,000", "Do not know", "Prefer not to say". We set the last two responses to NA and converted the remaining ones to levels 1-6, where 1 was the lowest salary band and 6 was the highest. |
| Age completed full time education | Participants were asked to indicate their educational qualifications in Field 6138. Those who did not indicate a university degree were asked when they stopped their full time education in Field 845. We used Field 845 and, those who indicated they did have a university degree were assigned (by us) an age of 22 for this variable, in order to keep them in the analysis. |
| Place of birth in the UK | Fields 129 (north coordinate) and 130 (east coordinate). Coordinates are given in metre-grid map units. |
| Home location | Fields 20074 (east coordinate) and 20075 (north coordinate). Coordinates are given in metre-grid map units. |

**Table S3.** Effect size estimates 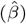 and strength of association (*p*-values) between PCs and scores as well as the association between the PCs and CAD in the white British subset of the UK Biobank. We provide here the 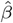 and *p*-values of each PC in a linear regression of the score (normalized within the white British subset) on the first three PCs calculated within the white British subset, age (defined using fields 32 and 34), sex (field 31), place of birth coordinates (fields 129 and 130), home location coordinates (fields 20074 and 20075), smoking status (field 20116), Townsend deprivation index at recruitment (field 189), income (field 738 converted into numeric values), and age when completed full time education (field 845), supplemented by qualifications (field 6138), where people who indicated they had a university or college degree were assumed to have completed their degree at 22). The results for CAD were from a logistic regression that used the same set of covariates. All continuous variables were scaled.

| Outcome | PC1 $\hat{\beta}$ ($p$-val) | PC2 $\hat{\beta}$ ($p$-val) | PC3 $\hat{\beta}$ ($p$-val) |
| --- | --- | --- | --- |
| metaGRS | -0.0038 (0.0555) | 0.0266 ($5.1 \times 10^{-42}$) | 0.0208 ($3.7 \times 10^{-32}$) |
| K2018 | 0.0135 ($1.1 \times 10^{-11}$) | 0.0048 (0.0145) | 0.0052 (0.0030) |
| E2020 | -0.0089 ($7.1 \times 10^{-6}$) | 0.0103 ($1.6 \times 10^{-7}$) | 0.0037 (0.0345) |
| CAD | -0.0158 (0.0740) | 0.0038 (0.6686) | 0.0147 (0.0593) |

**Table S4.**
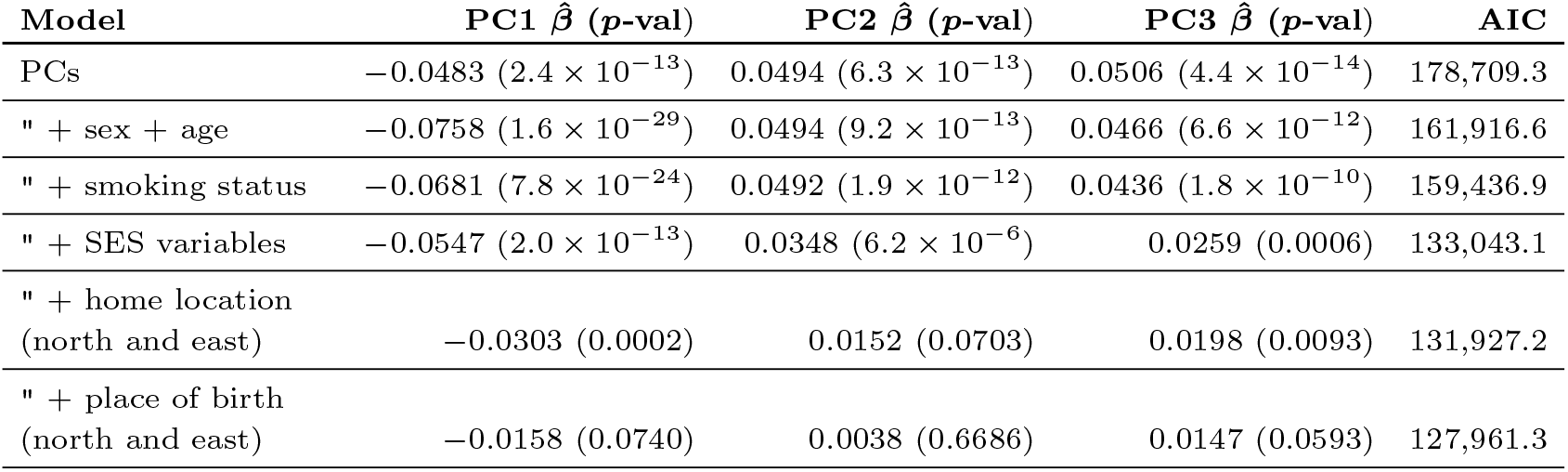
Effect size estimates and strength of association between PCs and CAD in the white British subset of the UK Biobank. We provide the effect size estimates 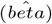, *p*-values, and the Akaike information criterion (AIC) for each model of each PC in a logistic regression of the score (normalized within the white British subset) on the first three PCs calculated within the 334,181 white British individuals for whom all the relevant covariates were available, sequentially adding environmental covariates to the model—that is, the model described in each row includes the covariates listed in the preceding rows as well. SES variables were the Townsend deprivation index, income, and age when completed full time education. All continuous variables were scaled.

| Model | PC1 $\hat{\beta}$ ($p$-val) | PC2 $\hat{\beta}$ ($p$-val) | PC3 $\hat{\beta}$ ($p$-val) | AIC |
| --- | --- | --- | --- | --- |
| PCs | -0.0483 ($2.4 \times 10^{-13}$) | 0.0494 ($6.3 \times 10^{-13}$) | 0.0506 ($4.4 \times 10^{-14}$) | 178,709.3 |
| " + sex + age | -0.0758 ($1.6 \times 10^{-29}$) | 0.0494 ($9.2 \times 10^{-13}$) | 0.0466 ($6.6 \times 10^{-12}$) | 161,916.6 |
| " + smoking status | -0.0681 ($7.8 \times 10^{-24}$) | 0.0492 ($1.9 \times 10^{-12}$) | 0.0436 ($1.8 \times 10^{-10}$) | 159,436.9 |
| " + SES variables | -0.0547 ($2.0 \times 10^{-13}$) | 0.0348 ($6.2 \times 10^{-6}$) | 0.0259 (0.0006) | 133,043.1 |
| " + home location<br>(north and east) | -0.0303 (0.0002) | 0.0152 (0.0703) | 0.0198 (0.0093) | 131,927.2 |
| " + place of birth<br>(north and east) | -0.0158 (0.0740) | 0.0038 (0.6686) | 0.0147 (0.0593) | 127,961.3 |

**Table S5.** Effect size estimates, *p*-values of association of each risk score with CAD, and CAD prevalence, calculated for each subpopulation. Scores were normalized within each subpopulation and included with age, sex, and the first 10 principal components provided by the UK Biobank.

| Population | Size | metaGRS $\hat{\beta}$ ($p$) | K2018 $\hat{\beta}$ ($p$) | E2020 $\hat{\beta}$ ($p$) | CAD prevalence |
| --- | --- | --- | --- | --- | --- |
| White British | 408,550 | 0.527 ($< 10^{-100}$) | 0.514 ($< 10^{-100}$) | 0.404 ($< 10^{-100}$) | 0.0572 |
| Irish | 12,668 | 0.544 ($1.1 \times 10^{-42}$) | 0.515 ($2.5 \times 10^{-39}$) | 0.430 ($3.5 \times 10^{-28}$) | 0.0603 |
| Other white | 16,208 | 0.555 ($2.3 \times 10^{-36}$) | 0.481 ($1.5 \times 10^{-30}$) | 0.310 ($2.4 \times 10^{-14}$) | 0.0436 |
| South Asian | 7,627 | 0.464 ($5.0 \times 10^{-27}$) | 0.405 ($1.2 \times 10^{-21}$) | 0.394 ($7.4 \times 10^{-21}$) | 0.0992 |
| Black, Afr., or<br>Carib. | 7,648 | 0.230 (0.001) | 0.187 (0.011) | 0.084 (0.250) | 0.0288 |
| Chinese | 1,502 | 0.319 (0.070) | 0.350 (0.047) | 0.313 (0.058) | 0.0262 |

**Table S6.**
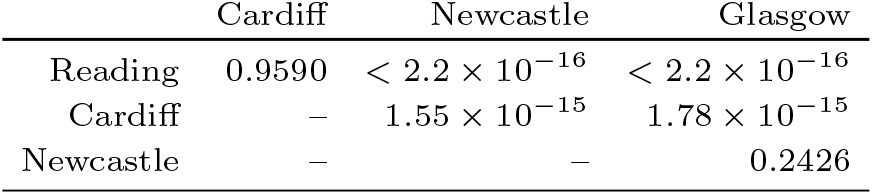
*P*-values of Kolmogorov-Smirnov tests of the pairwise equivalence of the distributions of the metaGRS scores calculated for white British individuals who attended one of the two indicated assessment centers. Low *p*-values indicate a rejection of the null hypothesis that the metaGRS score distributions are the same for the two assessment centers.

|  | Cardiff | Newcastle | Glasgow |
| --- | --- | --- | --- |
| Reading | 0.9590 | $< 2.2 \times 10^{-16}$ | $< 2.2 \times 10^{-16}$ |
| Cardiff | — | $1.55 \times 10^{-15}$ | $1.78 \times 10^{-15}$ |
| Newcastle | — | — | 0.2426 |

**Table S7.** *P*-values of Kolmogorov-Smirnov tests of the pairwise equivalence of the distributions of the K2018 scores calculated for white British individuals who attended one of the two indicated assessment centers. Low *p*-values indicate a rejection of the null hypothesis that the K2018 score distributions are the same for the two assessment centers.

|  | Cardiff | Newcastle | Glasgow |
| --- | --- | --- | --- |
| Reading | 0.5549 | 0.0004 | 0.9053 |
| Cardiff | – | 0.0075 | 0.9708 |
| Newcastle | – | – | 0.0126 |

**Table S8.** *P*-values of Kolmogorov-Smirnov tests of the pairwise equivalence of the distributions of the E2020 scores calculated for white British individuals who attended one of the two indicated assessment centers. Low *p*-values indicate a rejection of the null hypothesis that the E2020 score distributions are the same for the two assessment centers.

|  | Cardiff | Newcastle | Glasgow |
| --- | --- | --- | --- |
| Reading | 0.0485 | $7.17 \times 10^{-6}$ | $4.96 \times 10^{-6}$ |
| Cardiff | – | 0.0639 | 0.0003 |
| Newcastle | – | – | 0.0490 |

**Table S9.** Proportion of overlap among the white British individuals identified by each score as being in the top 10% of CAD risk (upper triangle) to the bottom 10% (lower triangle).

|  | metaGRS | K2018 | E2020 |
| --- | --- | --- | --- |
| metaGRS | – | 0.5530 | 0.3894 |
| K2018 | 0.5381 | – | 0.4520 |
| E2020 | 0.3906 | 0.4597 | – |

**Table S10.**
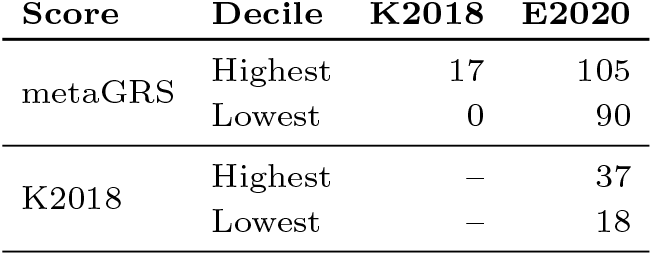
Number of people who are assigned by the score in the left-most column a level of risk (highest or lowest decile) who were placed in the opposite extreme risk category by the other two scores. For example, the first line tells us that there were 17 people who were in the highest decile of metaGRS scores who were also in the lowest decile of K2018 scores. Similarly, there were 105 people who were in the highest decile of metaGRS scores who were in the lowest decile of E2020 scores. The next line tells us the inverse: how many people were in the lowest decile of risk for the metaGRS who were also in the highest decile of risk for the other two scores. For reference, each decile contains 40,858 individuals.

**Fig. S1.**
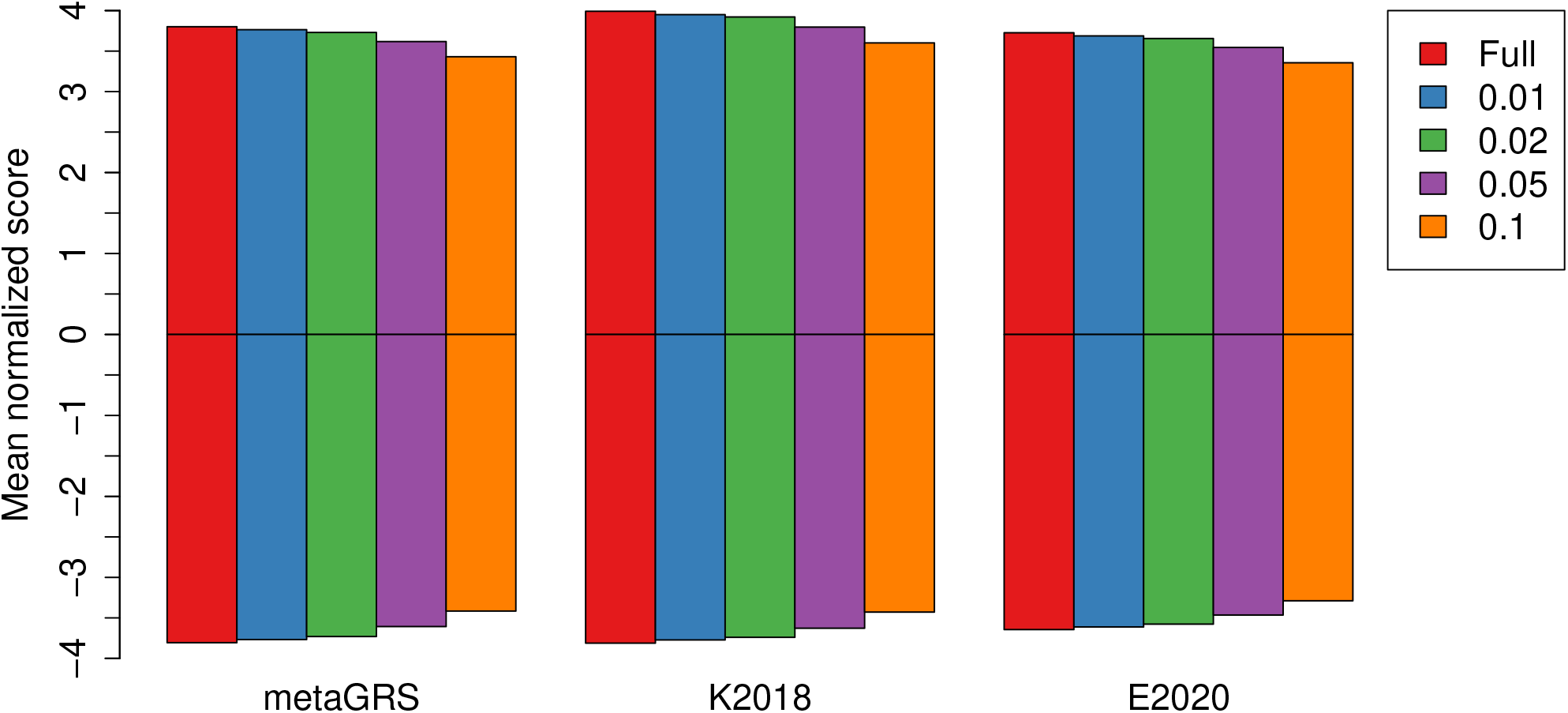
Mean normalized PRS for different rates of random missingness. The positive barplot shows the mean of the individuals who were identified in the original “full” score as having the highest 100 scores, and the negative shows the mean for those with the lowest 100 scores. The red bars show the values using the original “full” scores. Subsequent bars show the values of the scores after setting 1%, 2%, 5%, and 10% of the genotypes of these individuals at markers used in each score to missing.The distributions across the three scores are significantly different according to a Kolmogorov-Smirnov test.

## References

1. Naomi R. Wray, Tian Lin, Jehannine Austin, John J. McGrath, Ian B. Hickie, Graham K. Murray, and Peter M. Visscher. From Basic Science to Clinical Application of Polygenic Risk Scores: A Primer. JAMA Psychiatry, 78(1):101, January 2021.

2. Amit V. Khera, Mark Chaffin, Krishna G. Aragam, Mary E. Haas, Carolina Roselli, Seung Hoan Choi, Pradeep Natarajan, Eric S. Lander, Steven A. Lubitz, Patrick T. Ellinor, and Sekar Kathiresan. Genome-wide polygenic scores for common diseases identify individuals with risk equivalent to monogenic mutations. Nature Genetics, 50(9):1219–1224, September 2018. Number: 9.

3. Michael Inouye, Gad Abraham, Christopher P. Nelson, Angela M. Wood, Michael J. Sweeting, Frank Dudbridge, Florence Y. Lai, Stephen Kaptoge, Marta Brozynska, Tingting Wang, Shu Ye, Thomas R. Webb, Martin K. Rutter, Ioanna Tzoulaki, Riyaz S. Patel, Ruth J.F. Loos, Bernard Keavney, Harry Hemingway, John Thompson, Hugh Watkins, Panos Deloukas, Emanuele Di Angelantonio, Adam S. Butterworth, John Danesh, and Nilesh J. Samani. Genomic Risk Prediction of Coronary Artery Disease in 480,000 Adults. Journal of the American College of Cardiology, 72(16):1883–1893, October 2018. Number: 16.

4. Stephen Burgess and Simon G Thompson. Use of allele scores as instrumental variables for Mendelian randomization. International Journal of Epidemiology, 42(4):1134–1144, August 2013. Number: 4.

5. Karani S. Vimaleswaran, Diane J. Berry, Chen Lu, Emmi Tikkanen, Stefan Pilz, Linda T. Hiraki, Jason D. Cooper, Zari Dastani, Rui Li, and Denise K. Houston. Causal relationship between obesity and vitamin D status: bi-directional Mendelian randomization analysis of multiple cohorts. PLoS Med, 10(2):e1001383, 2013. Number: 2.

6. Ali Torkamani, Nathan E. Wineinger, and Eric J. Topol. The personal and clinical utility of polygenic risk scores. Nature Reviews Genetics, 19(9):581–590, September 2018. Number: 9.

7. Hannah Wand, Samuel A. Lambert, Cecelia Tamburro, Michael A. Iacocca Jack W. O’Sullivan, Catherine Sillari, Iftikhar J. Kullo, Robb Rowley, Jacqueline S. Dron, Deanna Brockman, Eric Venner, Mark I. McCarthy, Antonis C. Antoniou, Douglas F. Easton, Robert A. Hegele, Amit V. Khera, Nilanjan Chatterjee, Charles Kooperberg, Karen Edwards, Katherine Vlessis, Kim Kinnear, John N. Danesh, Helen Parkinson, Erin M. Ramos, Megan C. Roberts, Kelly E. Ormond, Muin J. Khoury, A. Cecile J. W. Janssens, Katrina A. B. Goddard, Peter Kraft, Jaqueline A. L. MacArthur, Michael Inouye, and Genevieve L. Wojcik. Improving reporting standards for polygenic scores in risk prediction studies. Nature, 591(7849):211–219, March 2021.

8. Simon Haworth, Ruth Mitchell, Laura Corbin, Kaitlin H. Wade, Tom Dudding, Ashley Budu-Aggrey, David Carslake, Gibran Hemani, Lavinia Paternoster, George Davey Smith, Neil Davies, Daniel J. Lawson, and Nicholas J. Timpson. Apparent latent structure within the UK Biobank sample has implications for epidemiological analysis. Nature Communications, 10(1):333, December 2019. Number: 1.

9. Arslan A Zaidi and Iain Mathieson. Demographic history mediates the effect of stratification on polygenic scores. eLife, 9:e61548, November 2020.

10. C. A. Anderson, F. H. Pettersson, G. M. Clarke, L. R. Cardon, A. P. Morris, and K. T. Zondervan. Data quality control in genetic case-control association studies. Nat Protoc, 5(9):1564–73, September 2010. Edition: 2010/11/19.

11. Joshua Elliott, Barbara Bodinier, Tom A. Bond, Marc Chadeau-Hyam, Evangelos Evangelou, Karel G. M. Moons, Abbas Dehghan, David C. Muller, Paul Elliott, and Ioanna Tzoulaki. Predictive Accuracy of a Polygenic Risk Score–Enhanced Prediction Model vs a Clinical Risk Score for Coronary Artery Disease. JAMA, 323(7):636, February 2020. Number: 7.

12. Bjarni J. Vilhjálmsson, Jian Yang, Hilary K. Finucane, Alexander Gusev, Sara Lindström, Stephan Ripke, Giulio Genovese, Po-Ru Loh, Gaurav Bhatia, Ron Do, Tristan Hayeck, Hong-Hee Won, Sekar Kathiresan, Michele Pato, Carlos Pato, Rulla Tamimi, Eli Stahl, Noah Zaitlen, Bogdan Pasaniuc, Gillian Belbin, Eimear E. Kenny, Mikkel H. Schierup, Philip De Jager, Nikolaos A. Patsopoulos, Steve McCarroll, Mark Daly, Shaun Purcell, Daniel Chasman, Benjamin Neale, Michael Goddard, Peter M. Visscher, Peter Kraft, Nick Patterson, Alkes L. Price, Stephan Ripke, Benjamin M. Neale, Aiden Corvin, James T.R. Walters, Kai-How Farh, Peter A. Holmans, Phil Lee, Brendan Bulik-Sullivan, David A. Collier, Hailiang Huang, Tune H. Pers, Ingrid Agartz, Esben Agerbo, Margot Albus, Madeline Alexander, Farooq Amin, Silviu A. Bacanu, Martin Begemann, Richard A. Belliveau, Judit Bene, Sarah E. Bergen, Elizabeth Bevilacqua, Tim B. Bigdeli, Donald W. Black, Richard Bruggeman, Nancy G. Buccola, Randy L. Buckner, William Byerley, Wiepke Cahn, Guiqing Cai, Dominique Campion, Rita M. Cantor, Vaughan J. Carr, Noa Carrera, Stanley V. Catts, Kimberly D. Chambert, Raymond C.K. Chan, Ronald Y.L. Chen, Eric Y.H. Chen, Wei Cheng, Eric F.C. Cheung, Siow Ann Chong, C. Robert Cloninger, David Cohen, Nadine Cohen, Paul Cormican, Nick Craddock, James J. Crowley, David Curtis, Michael Davidson, Kenneth L. Davis, Franziska Degenhardt, Jurgen Del Favero, Lynn E. DeLisi, Ditte Demontis, Dimitris Dikeos, Timothy Dinan, Srdjan Djurovic, Gary Donohoe, Elodie Drapeau, Jubao Duan, Frank Dudbridge, Naser Durmishi, Peter Eichhammer, Johan Eriksson, Valentina Escott-Price, Laurent Essioux, Ayman H. Fanous, Martilias S. Farrell, Josef Frank, Lude Franke, Robert Freedman, Nelson B. Freimer, Marion Friedl, Joseph I. Friedman, Menachem Fromer, Giulio Genovese, Lyudmila Georgieva, Elliot S. Gershon, Ina Giegling, Paola Giusti-Rodrguez, Stephanie Godard, Jacqueline I. Goldstein, Vera Golimbet, Srihari Gopal, Jacob Gratten, Jakob Grove, Lieuwe de Haan, Christian Hammer, Marian L. Hamshere, Mark Hansen, Thomas Hansen, Vahram Haroutunian, Annette M. Hartmann, Frans A. Henskens, Stefan Herms, Joel N. Hirschhorn, Per Hoffmann, Andrea Hofman, Mads V. Hollegaard, David M. Hougaard, Masashi Ikeda, Inge Joa, Antonio Julia, Rene S. Kahn, Luba Kalaydjieva, Sena Karachanak-Yankova, Juha Karjalainen, David Kavanagh, Matthew C. Keller, Brian J. Kelly, James L. Kennedy, Andrey Khrunin, Yunjung Kim, Janis Klovins, James A. Knowles, Bettina Konte, Vaidutis Kucinskas, Zita Ausrele Kucinskiene, Hana Kuzelova-Ptackova, Anna K. Kahler, Claudine Laurent, Jimmy Lee Chee Keong, S. Hong Lee, Sophie E. Legge, Bernard Lerer, Miaoxin Li, Tao Li, Kung-Yee Liang, Jeffrey Lieberman, Svetlana Limborska, Carmel M. Loughland, Jan Lubinski, Jouko Lnnqvist, Milan Macek, Patrik K.E. Magnusson, Brion S. Maher, Wolfgang Maier, Jacques Mallet, Sara Marsal ManuelMattheisen, Morten Robert W. McCarley,Colm Mattingsdal McDonald, Andrew M. McIntosh, Sandra Meier, Carin J. Meijer, Bela Melegh, Ingrid Melle, Raquelle I. Mesholam-Gately, Andres Metspalu, Patricia T. Michie, Lili Milani, Vihra Milanova, Younes Mokrab, Derek W. Morris, Ole Mors, Preben B. Mortensen, Kieran C. Murphy, Robin M. Murray, Inez Myin-Germeys, Bertram Mller-Myhsok, Mari Nelis, Igor Nenadic, Deborah A. Nertney, Gerald Nestadt, Kristin K. Nicodemus, Liene Nikitina-Zake, Laura Nisenbaum, Annelie Nordin, Eadbhard O’Callaghan, Colm O’Dushlaine, F. Anthony O’Neill, Sang-Yun Oh, Ann Olincy, Line Olsen, Jim Van Os, Christos Pantelis, George N. Papadimitriou, Sergi Papiol, Elena Parkhomenko, Michele T. Pato, Tiina Paunio, Milica Pejovic-Milovancevic, Diana O. Perkins, Olli Pietilinen, Jonathan Pimm, Andrew J. Pocklington, John Powell, Alkes Price, Ann E. Pulver, Shaun M. Purcell, Digby Quested, Henrik B. Rasmussen, Abraham Reichenberg, Mark A. Reimers, Alexander L. Richards, Joshua L. Roffman, Panos Roussos, Douglas M. Ruderfer, Veikko Salomaa, Alan R. Sanders, Ulrich Schall, Christian R. Schubert, Thomas G. Schulze, Sibylle G. Schwab, Edward M. Scolnick, Rodney J. Scott, Larry J. Seidman, Jianxin Shi, Engilbert Sigurdsson, Teimuraz Silagadze, Jeremy M. Silverman, Kang Sim, Petr Slominsky, Jordan W. Smoller, Hon-Cheong So, Chris C.A. Spencer, Eli A. Stahl, Hreinn Stefansson, Stacy Steinberg, Elisabeth Stogmann, Richard E. Straub, Eric Strengman, Jana Strohmaier, T. Scott Stroup, Mythily Subramaniam, Jaana Suvisaari, Dragan M. Svrakic, Jin P. Szatkiewicz, Erik Sderman, Srinivas Thirumalai, Draga Toncheva, Paul A. Tooney, Sarah Tosato, Juha Veijola, John Waddington, Dermot Walsh, Dai Wang, Qiang Wang, Bradley T. Webb, Mark Weiser, Dieter B. Wildenauer, Nigel M. Williams, Stephanie Williams, Stephanie H. Witt, Aaron R. Wolen, Emily H.M. Wong, Brandon K. Wormley, Jing Qin Wu, Hualin Simon Xi, Clement C. Zai, Xuebin Zheng, Fritz Zimprich, Naomi R. Wray, Kari Stefansson, Peter M. Visscher, Rolf Adolfsson, Ole A. Andreassen, Douglas H.R. Blackwood, Elvira Bramon, Joseph D. Buxbaum Anders D. Børglum, Sven Cichon, Ariel Darvasi, Enrico Domenici, Hannelore Ehrenreich, Tonu Esko, Pablo V. Gejman, Michael Gill, Hugh Gurling, Christina M. Hultman, Nakao Iwata, Assen V. Jablensky, Erik G. Jonsson, Kenneth S. Kendler, George Kirov, Jo Knight, Todd Lencz, Douglas F. Levinson, Qingqin S. Li, Jianjun Liu, Anil K. Malhotra, Steven A. McCarroll, Andrew McQuillin, Jennifer L. Moran, Preben B. Mortensen, Bryan J. Mowry, Markus M. Nthen, Roel A. Ophoff, Michael J. Owen, Aarno Palotie, Carlos N. Pato, Tracey L. Petryshen, Danielle Posthuma, Marcella Rietschel, Brien P. Riley, Dan Rujescu, Pak C. Sham, Pamela Sklar, David St. Clair, Daniel R. Weinberger, Jens R. Wendland, Thomas Werge, Mark J. Daly, Patrick F. Sullivan Michael C. O’Donovan, Peter Kraft, David J. Hunter, Muriel Adank, Habibul Ahsan, Kristiina Aittomäki, Laura Baglietto, Sonja Berndt, Carl Blomquist, Federico Canzian, Jenny Chang-Claude, Stephen J. Chanock, Laura Crisponi, Kamila Czene, Norbert Dahmen, Isabel dos Santos Silva, Douglas Easton, A. Heather Eliassen, Jonine Figueroa, Olivia Fletcher, Montserrat Garcia-Closas, Mia M. Gaudet, Lorna Gibson, Christopher A. Haiman, Per Hall, Aditi Hazra, Rebecca Hein, Brian E. Henderson, Albert Hofman, John L. Hopper, Astrid Irwanto, Mattias Johansson, Rudolf Kaaks, Muhammad G. Kibriya, Peter Lichtner, Sara Lindström, Jianjun Liu, Eiliv Lund, Enes Makalic, Alfons Meindl, Hanne Meijers-Heijboer, Bertram Müller-Myhsok, Taru A. Muranen, Heli Nevanlinna, Petra H. Peeters, Julian Peto, Ross L. Prentice, Nazneen Rahman, María José Sánchez, Daniel F. Schmidt, Rita K. Schmutzler, Melissa C. Southey, Rulla Tamimi, Ruth Travis, Clare Turnbull, Andre G. Uitterlinden, Rob B. van der Luijt, Quinten Waisfisz, Zhaoming Wang, Alice S. Whittemore, Rose Yang, and Wei Zheng. Modeling Linkage Disequilibrium Increases Accuracy of Polygenic Risk Scores. The American Journal of Human Genetics, 97(4):576–592, October 2015. Number: 4.

13. Timothy Shin Heng Mak, Robert Milan Porsch, Shing Wan Choi, Xueya Zhou, and Pak Chung Sham. Polygenic scores via penalized regression on summary statistics: MAK et al. Genetic Epidemiology, 41(6):469–480, September 2017.

14. Clare Bycroft, Colin Freeman, Desislava Petkova, Gavin Band, Lloyd T. Elliott, Kevin Sharp, Allan Motyer, Damjan Vukcevic, Olivier Delaneau, Jared O’Connell, Adrian Cortes, Samantha Welsh, Alan Young, Mark Effingham, Gil McVean, Stephen Leslie, Naomi Allen, Peter Donnelly, and Jonathan Marchini. The UK Biobank resource with deep phenotyping and genomic data. Nature, 562(7726):203–209, October 2018. Number: 7726.

15. W. B. Provine. The role of mathematical population geneticists in the evolutionary synthesis of the 1930s and 1940s. Studies in History of Biology, 2:167–192, 1978.

16. Alkes L Price, Nick J Patterson, Robert M Plenge, Michael E Weinblatt, Nancy A Shadick, and David Reich. Principal components analysis corrects for stratification in genome-wide association studies. Nature Genetics, 38(8):904–909, August 2006. Number: 8.

17. Matthew L Freedman, David Reich, Kathryn L Penney, Gavin J McDonald, Andre A Mignault, Nick Patterson, Stacey B Gabriel, Eric J Topol, Jordan W Smoller, Carlos N Pato, Michele T Pato, Tracey L Petryshen, Laurence N Kolonel, Eric S Lander, Pamela Sklar, Brian Henderson, Joel N Hirschhorn, and David Altshuler. Assessing the impact of population stratification on genetic association studies. Nature Genetics, 36(4):388–393, April 2004. Number: 4.

18. Jonathan Marchini, Lon R Cardon, Michael S Phillips, and Peter Donnelly. The effects of human population structure on large genetic association studies. Nature Genetics, 36(5):512–517, May 2004. Number: 5.

19. Joel N. Hirschhorn and Mark J. Daly. Genomewide association studies for common diseases and complex traits. Nature Reviews Genetics, 6(2):95–108, February 2005. Number: 2.

20. Samsiddhi Bhattacharjee, Zhaoming Wang, Julia Ciampa, Peter Kraft, Stephen Chanock, Kai Yu, and Nilanjan Chatterjee. Using Principal Components of Genetic Variation for Robust and Powerful Detection of Gene-Gene Interactions in Case-Control and Case-Only Studies. The American Journal of Human Genetics, 86(3):331–342, March 2010. Number: 3.

21. David Reich, Alkes L. Price, and Nick Patterson. Principal component analysis of genetic data. Nature Genetics, 40(5):491–492, 2008. Number: 5.

22. Alicia R. Martin, Christopher R. Gignoux, Raymond K. Walters, Genevieve L. Wojcik, Benjamin M. Neale, Simon Gravel, Mark J. Daly, Carlos D. Bustamante, and Eimear E. Kenny. Human Demographic History Impacts Genetic Risk Prediction across Diverse Populations. The American Journal of Human Genetics, 100(4):635–649, April 2017. Number: 4.

23. Alicia R. Martin, Masahiro Kanai, Yoichiro Kamatani, Yukinori Okada, Benjamin M. Neale, and Mark J. Daly. Clinical use of current polygenic risk scores may exacerbate health disparities. Nature Genetics, 51(4):584–591, April 2019.

24. Hakhamanesh Mostafavi, Arbel Harpak, Ipsita Agarwal, Dalton Conley, Jonathan K Pritchard, and Molly Przeworski. Variable prediction accuracy of polygenic scores within an ancestry group. eLife, 9:e48376, January 2020.

25. Deo Mujwara, Geoffrey Henno, Stephen T. Vernon, Siyang Peng, Paolo Di Domenico, Brock Schroeder, George B. Busby, Gemma A Figtree, and Giordano Botta. Integrating a Polygenic Risk Score for Coronary Artery Disease as a Risk-Enhancing Factor in the Pooled Cohort Equation: A Cost-Effectiveness Analysis Study. Journal of the American Heart Association, 11(12):e025236, June 2022.

26. Mark Simcoe, Ana Valdes, Fan Liu, Nicholas A. Furlotte, David M. Evans, Gibran Hemani, Susan M. Ring, George Davey Smith, David L. Duffy, Gu Zhu, Scott D. Gordon, Sarah E. Medland, Dragana Vuckovic, Giorgia Girotto, Cinzia Sala, Eulalia Catamo, Maria Pina Concas, Marco Brumat, Paolo Gasparini, Daniela Toniolo, Massimiliano Cocca, Antonietta Robino, Seyhan Yazar, Alex Hewitt, Wenting Wu, Peter Kraft, Christopher J. Hammond, Yuan Shi, Yan Chen, Changqing Zeng, Caroline C. W. Klaver, Andre G. Uitterlinden, M. Arfan Ikram, Merel A. Hamer, Cornelia M. van Duijn, Tamar Nijsten, Jiali Han, David A. Mackey, Nicholas G. Martin, Ching-Yu Cheng, the 23andMe Research Team, the International Visible Trait Genetics Consortium, David A. Hinds, Timothy D. Spector, Manfred Kayser, and Pirro G. Hysi. Genome-wide association study in almost 195,000 individuals identifies 50 previously unidentified genetic loci for eye color. Science Advances, 7(11):eabd1239, March 2021.

27. Amit V. Khera, Connor A. Emdin, Isabel Drake, Pradeep Natarajan, Alexander G. Bick, Nancy R. Cook, Daniel I. Chasman, Usman Baber, Roxana Mehran, Daniel J. Rader, Valentin Fuster, Eric Boerwinkle, Olle Melander, Marju Orho-Melander, Paul M Ridker, and Sekar Kathiresan. Genetic risk, adherence to a healthy lifestyle, and coronary disease. New England Journal of Medicine, 375(24):2349–2358, 2016. PMID: 27959714.

28. A Cecile J W Janssens. Validity of polygenic risk scores: are we measuring what we think we are? Human Molecular Genetics, 28(R2):R143–R150, November 2019.

29. R Core Team. R: A Language and Environment for Statistical Computing. R Foundation for Statistical Computing, Vienna, Austria, 2020.

30. Ada Hui, Asam Latif, Kathryn Hinsliff-Smith, and Timothy Chen. Exploring the impacts of organisational structure, policy and practice on the health inequalities of marginalised communities: Illustrative cases from the UK healthcare system. Health Policy, 124(3):298–302, March 2020. Number: 3.

31. Gad Abraham, Yixuan Qiu, and Michael Inouye. FlashPCA2: principal component analysis of Biobank-scale genotype datasets. Bioinformatics, 33(17):2776–2778, September 2017. Number: 17.

32. Christopher C Chang, Carson C Chow, Laurent CAM Tellier, Shashaank Vattikuti, Shaun M Purcell, and James J Lee. Second-generation PLINK: rising to the challenge of larger and richer datasets. GigaScience, 4(1):7, December 2015. Number: 1.

33. Shaun Purcell and Christopher Chang. Plink 1.9b 5.2, 2019. Programmers: :n17703.

34. Amit V. Khera, Mark Chaffin, Kaitlin H. Wade, Sohail Zahid, Joseph Brancale, Rui Xia, Marina Distefano, Ozlem Senol-Cosar, Mary E. Haas, Alexander Bick, Krishna G. Aragam, Eric S. Lander, George Davey Smith, Heather Mason-Suares, Myriam Fornage, Matthew Lebo, Nicholas J. Timpson, Lee M. Kaplan, and Sekar Kathiresan. Polygenic Prediction of Weight and Obesity Trajectories from Birth to Adulthood. Cell, 177(3):587–596.e9, April 2019. Number: 3.

35. Samuel A. Lambert, Laurent Gil, Simon Jupp, Michael Chapman, Helen Parkinson, John Danesh, Jacqueline A.L. MacArthur, and Michael Inouye. The Polygenic Score (PGS) Catalog: an open database to enable reproducibility and systematic evaluation.

